# Geographic risk assessment of *Batrachochytrium salamandrivorans* invasion in Costa Rica as a means of informing emergence management and mitigation

**DOI:** 10.1101/2023.10.20.563237

**Authors:** Henry C. Adams, Katherine E. Markham, Marguerite Madden, Matthew J. Gray, Federico Bolanos Vives, Gerardo Chaves, Sonia M. Hernandez

## Abstract

Remotely-sensed risk assessments of emerging, invasive pathogens are key to targeted surveillance and outbreak responses. The recent emergence and spread of the fungal pathogen, *Batrachochytrium salamandrivorans* (*Bsal*), in Europe has negatively impacted multiple salamander species. Scholars and practitioners are increasingly concerned about the potential consequences of this lethal pathogen in the Americas, where salamander biodiversity is higher than anywhere else in the world. Although *Bsal* has not yet been detected in the Americas, certain countries have already proactively implemented monitoring and detection plans in order to identify areas of greatest concern and enable efficient contingency planning in the event of pathogen detection. To predict areas in Costa Rica with a high *Bsal* transmission risk, we employed ecological niche modeling combined with biodiversity and tourist visitation data to ascertain the specific risk to a country with world renowned biodiversity. Our findings indicate that approximately 23% of Costa Rica’s landmass provides suitable conditions for *Bsal*, posing a threat to 37 salamander species. The Central and Talamanca mountain ranges, in particular, have habitats predicted to be highly suitable for the pathogen. To facilitate monitoring and mitigation efforts, we identified eight specific protected areas that we believe are at the greatest risk due to a combination of high biodiversity, tourist visitation, and suitable habitat for *Bsal*. We advise regular monitoring utilizing remotely-sensed data and ecological niche modeling to effectively target *in-situ* surveillance and as places begin implementing educational efforts.

## Introduction

Emergent infectious pathogens and their associated diseases pose an increasing threat to global wildlife biodiversity and ecosystem health (1). This threat calls for proactive management strategies that utilize pathogen invasion risk predictive models, targeted surveillance, and susceptibility assessment. Such strategies can better mitigate negative outcomes, reduce costs, and promote more efficient response (2–4). In the case of chytridiomycosis caused by the fungal pathogen *Batrachochytrium dendrobatidis* (*Bd*), decades passed between initial emergence, identifying and describing its etiology, and subsequent unified action. It initially emerged in the 1970s and caused extensive amphibian declines and extinctions in many Central American and Australian species before its formal scientific description in 1998 (5–8). Despite that some regions experienced upwards of 40% *Bd* related amphibian biodiversity loss, it was not until 2005-2006 that Australia and the United States developed conservation action plans to combat the fungus (2). *Bd* has been implicated in the decline of over 500 amphibian species worldwide. Fortunately, observations of expanding remnant populations of *Bd*-impacted species and persistent research investigating the efficacy of species reintroductions and other management strategies shine hope on this conservation crisis (3,6,9,10).

Potentially, as a result of lessons learned from the response to *Bd*’s emergence, a more rapid response was mobilized with the emergence of *Pseudogymnoascus destructans* (*Pd*), the causative agent of White-Nose Syndrome in bats. Management action plans aimed to monitor and predict transmission and dispersal of the pathogen were drafted within two years of its description in 2008 in Howes Cave near Albany, New York (11–13). Such proactive responses have allowed for the relatively rapid development of national and innovative *Pd* surveillance in North America, management strategies that vary based on ecological variations (i.e., species, hibernation location, etc.), and anti-fungal treatment assessment and implementation (14–16).

Rapid proactive approaches to wildlife health management are crucial, especially in light of the recent emergence of the fungal pathogen, *Batrachochytrium salamandrivorans* (*Bsal*), the third chytrid fungus known to parasitize vertebrate hosts. *Bsal* is a saprophytic fungus that, while having an affinity for cool, moist environments, can persist at a fairly wide thermal range (∼5°- 25°C). The fungus, aptly dubbed the “salamander plague”, is most pathogenic between 10°-15°C and presents a major global threat to urodelans (17–22). *Bsal* originated in East Asia and was likely dispersed through the international pet trade (7,18,23–25). *Bsal* was first described by Martell and colleagues in 2013, after being isolated from impacted European fire salamanders (*Salamandra salamandra*) in the Netherlands. *Bsal*-associated mortality events had been documented in the Netherlands starting in 2008, with population declines upwards of 94% (17), although more recent studies have shown that *Bsal* emerged in Germany as early as 2004 (21,22).

The pathogen’s emergence in Europe was met with highly proactive research and management measures. Martel and colleagues initiated wide scale surveillance, surveying over 5,000 individual animals, and conducted experimental susceptibility infection trials to determine the pathogen’s significance to a large variety of hosts prior to its widespread emergence (18). In response to *Bsal’s* emergence, by 2016, both the European Union and the United States had enacted regulations limiting the international trade of salamander species (2,26). Further, susceptibility trials have continued to illuminate the nuances of *Bsal* susceptibility in salamanders and the ability of anurans to persist as subclinical, infectious hosts (19,20,25,27). Ongoing surveillance monitoring in Europe has documented the expansion of *Bsal’s* range, which has now been detected in wild salamander populations in Belgium, Germany, and Spain (3,18,23,28–32). Detections in Spain have been separated by over 1,000 km from previous reports, potentially highlighting the density independence of *Bsal* and the importance of continuous surveillance efforts (3,33,34). The detection was met with immediate collaborative efforts, including temporary wetland drainage and removal of *Bsal* positive individuals, between regional authorities and scientists that led to the containment of the pathogen (3).

The Americas are home to the world’s most diverse salamander communities (35). Although *Bsal* has yet to be detected in the Western hemisphere, the execution and expansion of proactive conservation measures, similar to those taken in Spain, are of immense importance (31,36). Specifically, Costa Rica possesses the fifth most diverse salamander community in the world as well as a wealth of habitats potentially ecologically suitable for *Bsal* persistence (17,19,24,37,38). Additionally, Costa Rica has a thriving ecotourism industry with the majority of tourists originating from Europe and the United States annually (14% (∼280,000) and 40% (∼800,000) respectively) (39). Fomites, such as outdoor recreational and research equipment, can facilitate the transmission of *Bd* and *Bsal* to naïve environments. Ecotourism and other anthropogenic activities exacerbated the emergence of *Bd* in Costa Rica during the 1980s and 1990s (7,40), and a similar narrative could unfold with *Bsal* in the absence of proper biosafety and responsible recreation. To date, only limited surveillance for *Bsal* has been conducted in Costa Rica, and it has failed to detect it in native salamander species in small geographic areas in the Cordillera Talamanca and Central (38).

Ecological Niche Models (ENM, also referred to as Species Distribution Models) are valuable tools for understanding species’ habitat and therefore generating efficient and targeted surveillance efforts. ENM have been employed to evaluate the risk of invasion of *Bd* in Costa Rica, and *Bsal* in North America, Europe, and Central and South America (9,29,35,41–43). García-Rodríguez and colleagues recently generated *Bsal* ENM for Central and South America that identified multiple hotspots for *Bsal* introduction, mostly located in central and southern Mexico and Costa Rican mountain ranges (42,43). Despite the fact that the work by García-Rodríguez et al. (43) identifies sites where the diversity of salamanders and an optimal environment for *Bsal* converge, we believe that, in order to have a preventive effect on the transmission of the pathogen, we must consider the most probable sources of transmission and their relationship with the divers and susceptible areas. Indeed, recent work modeling *Bsal* spread in Europe has found that trail density is an important predictor of the pathogen’s presence (44). Our work advances our understanding of *Bsal* risk by modeling potential habitat using a smaller geographic extent than considered previously, potentially providing a more regionally specific risk assessment that accurately predicts occurrence (45) although see (46).

We sought to identify areas of high risk in Costa Rica based on 1) *Bsal* ecological suitability, 2) species diversity of all urodelan, and 3) the level of human visitation as a continuous source of possible pathogens. Here, we developed an ENM similar to García-Rodríguez et al. (43) to identify areas ecologically suitable for *Bsal* and used data provided by the International Union for Conservation of Nature (IUCN) and the Costa Rican Institute of Tourism (abbreviation here) to evaluate amphibian distributions and human visitations, respectively. Our findings have the potential to provide a more nuanced spatial analysis of potential risk at a higher spatial resolution, leading to more precise and targeted monitoring efforts.

## Methods

To identify areas in Costa Rica at high risk of *Bsal* introduction, we considered amphibian alpha biodiversity and habitat suitability to *Bsal* based on its current natural range (18,24,47). Areas that are both highly suitable for *Bsal* and have high amphibian biodiversity are identified as high priority monitoring areas. All modeling was done using TerrSet’s Habitat and Biodiversity Modeler (48).

Alpha diversity, or the total number of species within an area (roughly one km^2^), was calculated using species’ extent data from the IUCN Red List (49,50). Alpha diversity is mapped on a continuous scale. To more easily communicate findings, we reclassified salamander alpha diversity into four qualitative categories: low (having no or one species of salamanders), medium (two or three salamander sp.), high (four or five), and very high (six to eight). Because of their potential to act as infectious, sub-clinical hosts for *Bsal*, anuran alpha diversity was also calculated and reclassified into four qualitative categories: low (eight or fewer species), medium (nine to 26 species), high (27 to 39 species), and very high (over 39 species) (19,25). As monitoring and mitigation efforts are likely to be more feasible inside protected areas, we also calculated the gamma diversity, or species richness, of salamander species across all protected areas. Gamma diversity is useful when comparing an ecosystem or region’s relative diversity to prioritize areas for conservation. The Range Restriction Index (RRI), which compares the area a species inhabits with the entire study region (Costa Rica), was also calculated for salamanders. Higher RRI values suggest that the majority of species present have relatively restricted ranges and are highly endemic.

### Modeling potential Bsal habitat

To identify potential suitable habitat for *Bsal* in Costa Rica, ecological niche modeling was conducted using *Bsal* occurrence records from within its endemic range. We used the maximum entropy model (Maxent), a correlative model that relates known species occurrences with environmental variables (51). Maxent models distribution using presence-only data is highly accurate (52) and performs well with smaller sample sizes (53). Selecting only occurrence data from regions where *Bsal* is endemic, we trained our model using 34 presence points from previously published sources (24,47,54). Environmental variables included in modeling *Bsal* habitat were mean diurnal range, maximum temperature of the warmest month, temperature annual range, precipitation seasonality, precipitation of warmest quarter, and precipitation of coldest quarter (42,55). These environmental variables align with previous studies predicting *Bsal* suitability (42). Correlation between environmental variables was checked, and all variables scored below 0.7 (56,57).

To predict species occurrence in Costa Rica, we used a parametrization of regularization multiplier (the degree to which predictions are fitted to training data) of 2.5 to obtain a more generalized output i.e., predictions are not overfitted to training data. We also used linear, quadratic, and product feature class combinations so that the mean of each environmental variable in predicted occurrences matched that of observed occurrences, the variance in environmental variables for the predicted occurrences is constrained to that of observed occurrences, and the covariance in environmental variables is constrained with other predictor environmental variables. Maxent produces a map ranging from zero to one, with one indicating the most highly suitable habitat. We reclassified the suitability map into low (less than 0.5), medium (0.5-0.75), and high (over 0.75) suitability categories to simplify the interpretation of results.

High priority protected areas where monitoring efforts should be focused were identified as such if 1) *Bsal* suitability was high and caudata alpha diversity was high/very high or 2) *Bsal* suitability was moderate, Caudata alpha diversity was high/very high, and reported annual visitation was above 40,000 individuals (the top 25% of human visitation for protected areas). We also compared these high priority areas with the RRI to identify species with particularly restricted ranges (i.e., at higher risk of local extinctions). An overview of the overall process for identifying *Bsal* monitoring areas in Costa Rica is shown in Figure 1.

**Figure 1.**
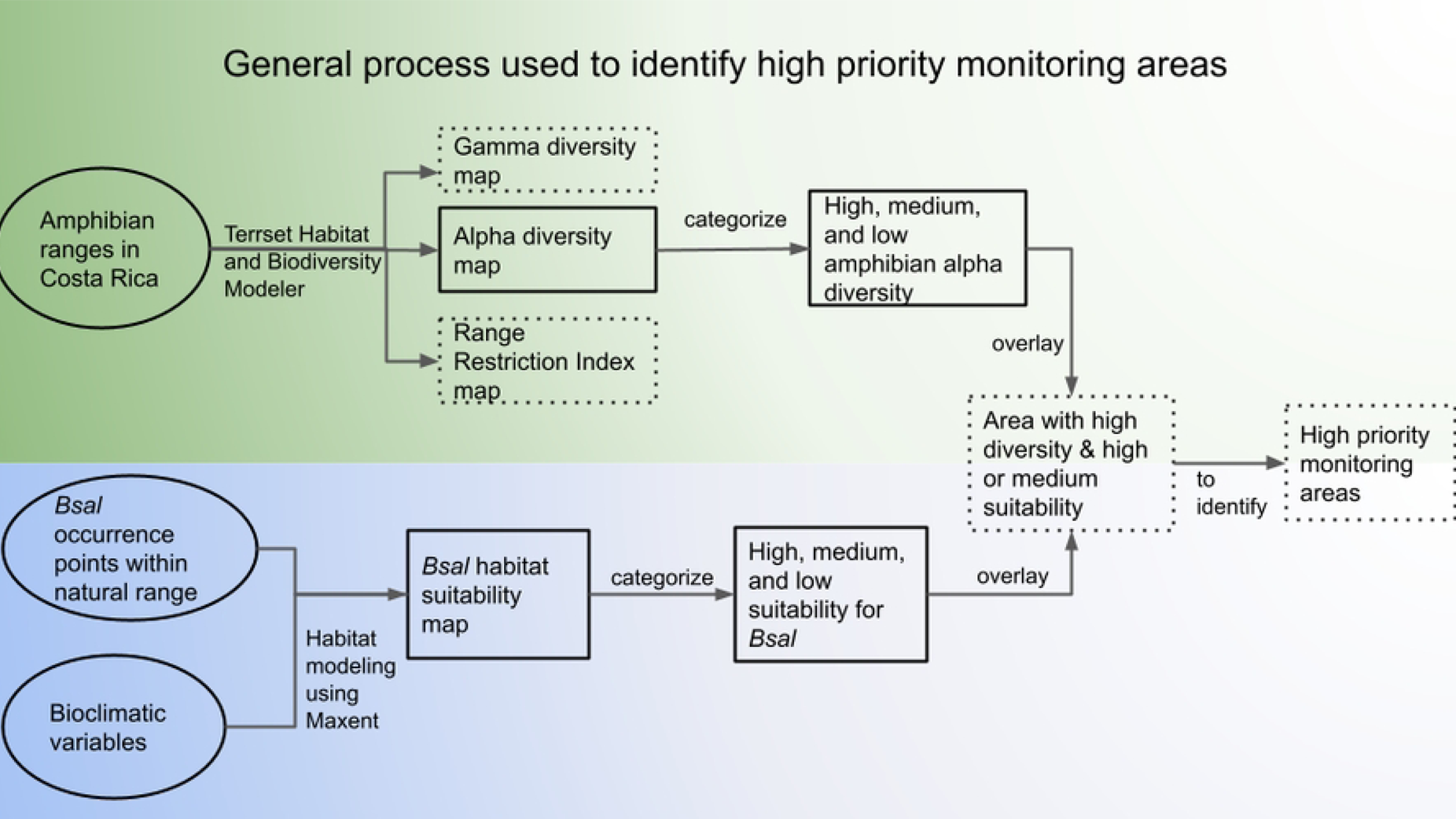
Process of identifying high priority monitoring areas. Conceptual diagram showing the data, tools, and general processes used to identify areas highly suitable for *Bsal* that also have high levels of amphibian diversity. Circles indicate input data, outputs are outlined by squares, and final outputs are outlined by square dotted lines. study methods Components of biodiversity modeling using amphibian data shown with green gradient and components of habitat suitability mapping of *Bsal* shown with blue gradient.

In addition to amphibian biodiversity and *Bsal* habitat suitability, we examined tourist visitation to protected areas using data provided by the Costa Rican Ecotourism Institute. Visitation data from 2018 were available for 46 out of 164 protected areas and included both national and international tourists (Figure 2). A list of the parks for which visitation data was available can be found in Table I, Appendix I. Protected area data is from Protected Planet (58) and includes national parks, biological reserves, and United Nations Educational, Scientific and Cultural Organization (UNESCO) World Heritage Sites. By finding where high tourist visitation in parks overlaps with high/very high alpha diversity and high or moderate *Bsal* suitability, we further identified areas of high priority for pathogen surveillance and mitigative strategies.

**Figure 2.**
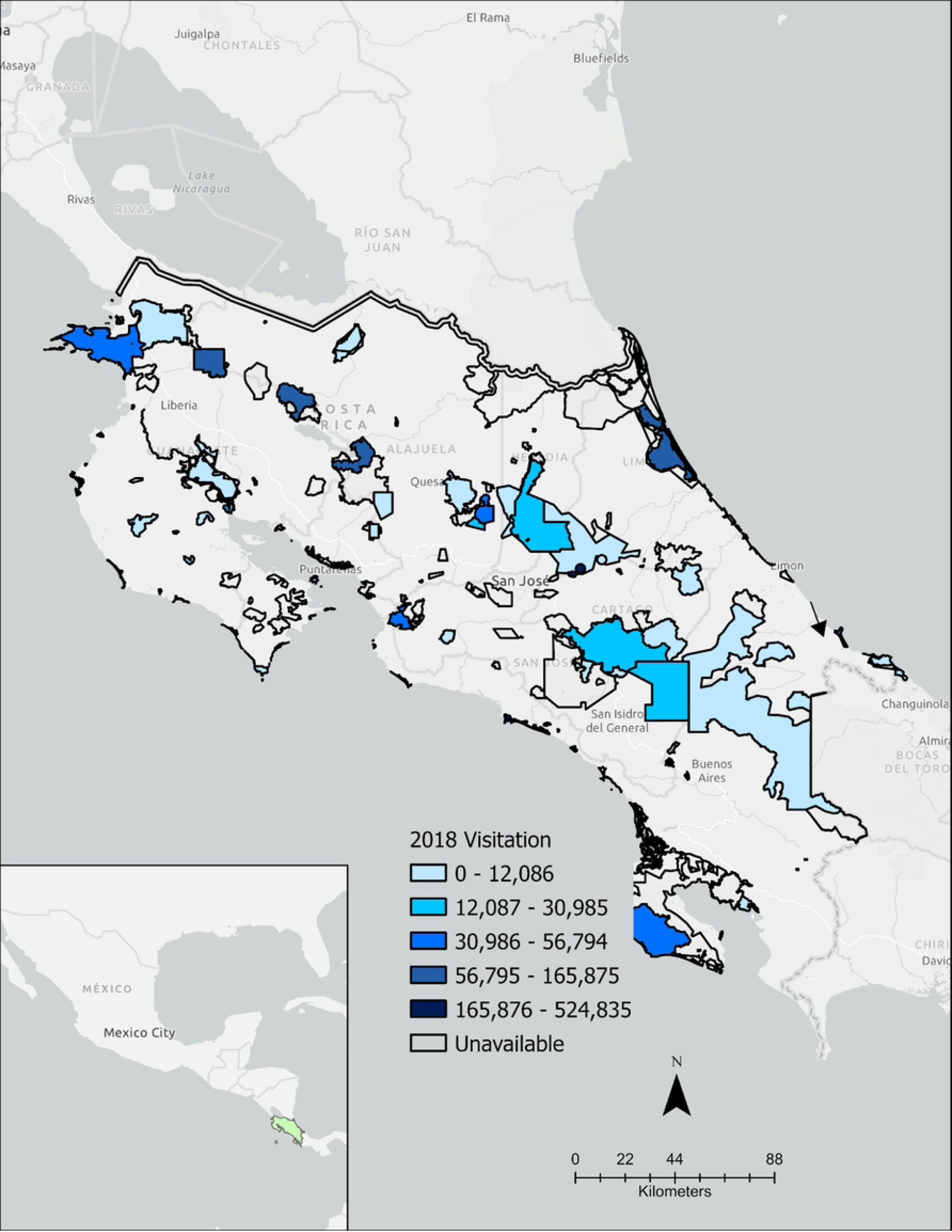
2018 Human visitation in Costa Rica protected areas. Protected areas include wildlife refuges, biological reserves, national parks, and world heritage sites. Protected areas are outlined in black. Where available, visitation numbers within protected areas for 2018 are indicated by color, with darker blue indicating more visitors. Insert map shows the geographic location of Costa Rica as a land bridge between North and South America.

## Results

### Diversity results

Alpha diversity for salamanders was greatest in the central part of Costa Rica as well as along the Panamanian border and lowest in northwestern Costa Rica (Figure 3). Protected areas with high amphibian gamma diversity, or total biodiversity for Costa Rica, include Parque Internacional La Amistad, Parque Nacional Braulio Carrillo, Parque Nacional Tapantí - Macizo Cerro de la Muerte, Reserva Forestal Río Macho, and Reserva de la Biosfera Cordillera Volcánica Central (Figure III in Appendix). Anuran alpha diversity followed similar patterns, with high diversity along the eastern portion of the country overlapping areas of high salamander diversity.

**Figure 3.**
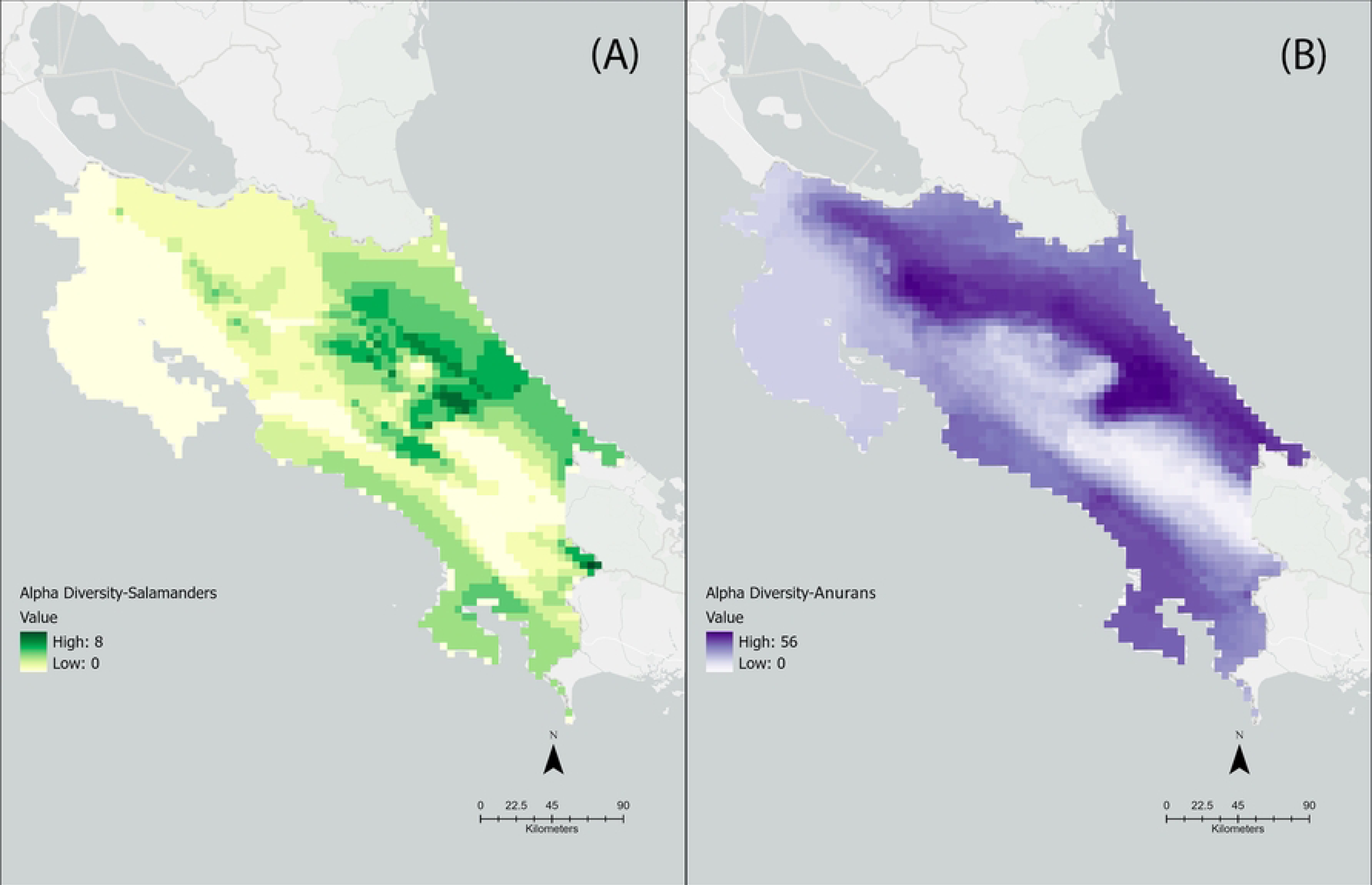
Costa Rican salamander and anuran alpha diversity. These maps denote alpha diversity of salamanders (A) and anurans (B) in Costa Rica. Darker shades indicate higher alpha diversity. Alpha diversity was calculated by summing the total number of salamander species in each ∼1 km by 1 km pixel.

### General Bsal results

Based on our results, the Talamanca Mountain Range, the Central Mountain Range, and the Northern Caribbean coast of Costa Rica have the most suitable habitat conditions for *Bsal* (Figure 4). *Bsal* modeled habitat suitability scores range from zero (unsuitable) to one (highly suitable), and the highest scores observed approached 0.80. Thus, of Costa Rica’s roughly 51,000 km^2^ landmass, we classified 77.33% as low (scores <0.50), 22.72% as medium (scores 0.5-0.75), and 0.15% as high suitability (scores >0.75) for *Bsal* (Figure 5).

**Figure 4.**
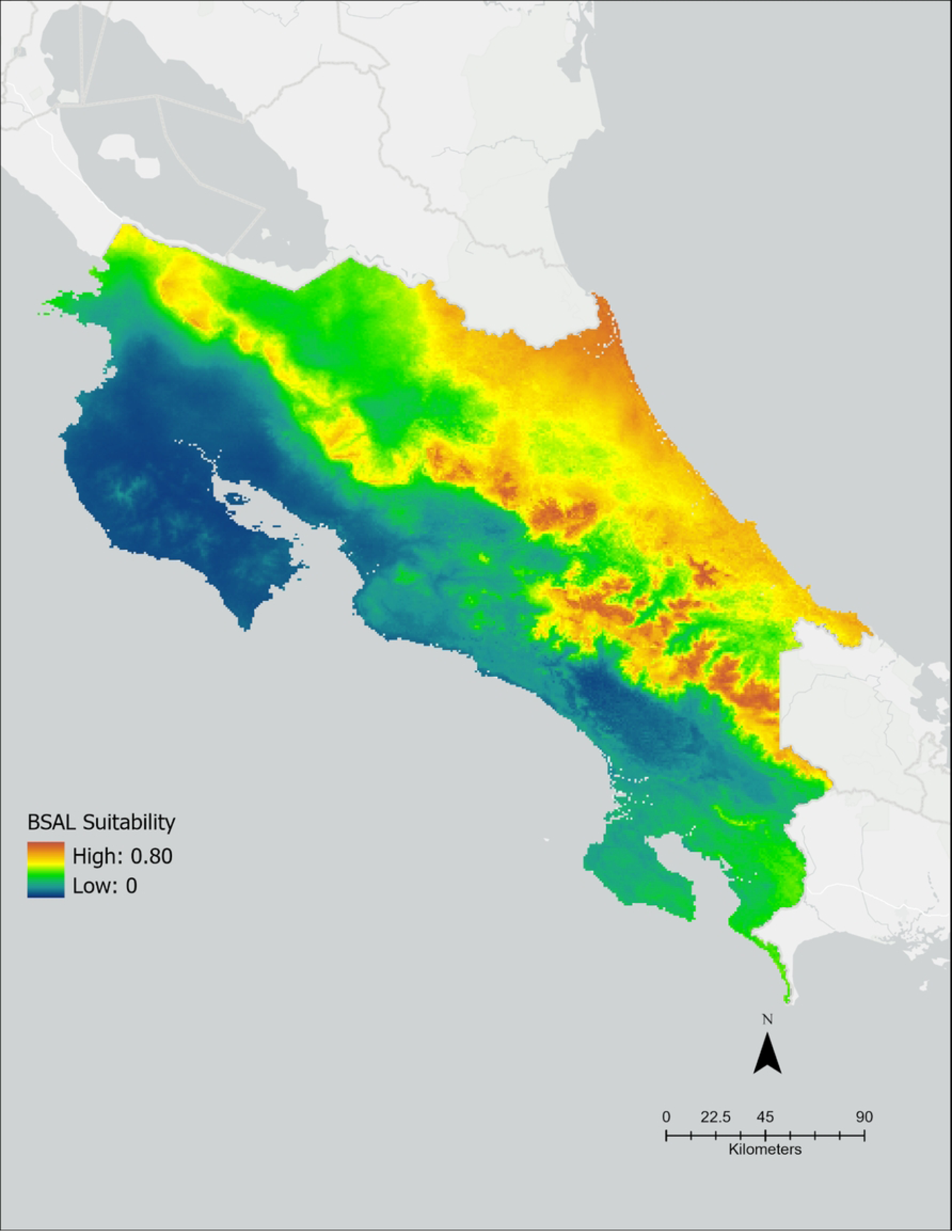
Predicted *Bsal* suitability in Costa Rica. Higher suitability is indicated by reds and oranges and can be observed in the Cordillera Central and Cordillera Talamanca mountain ranges, as well as along the northeastern Caribbean slope. Lower suitability is indicated by blues and greens.

**Figure 5.**
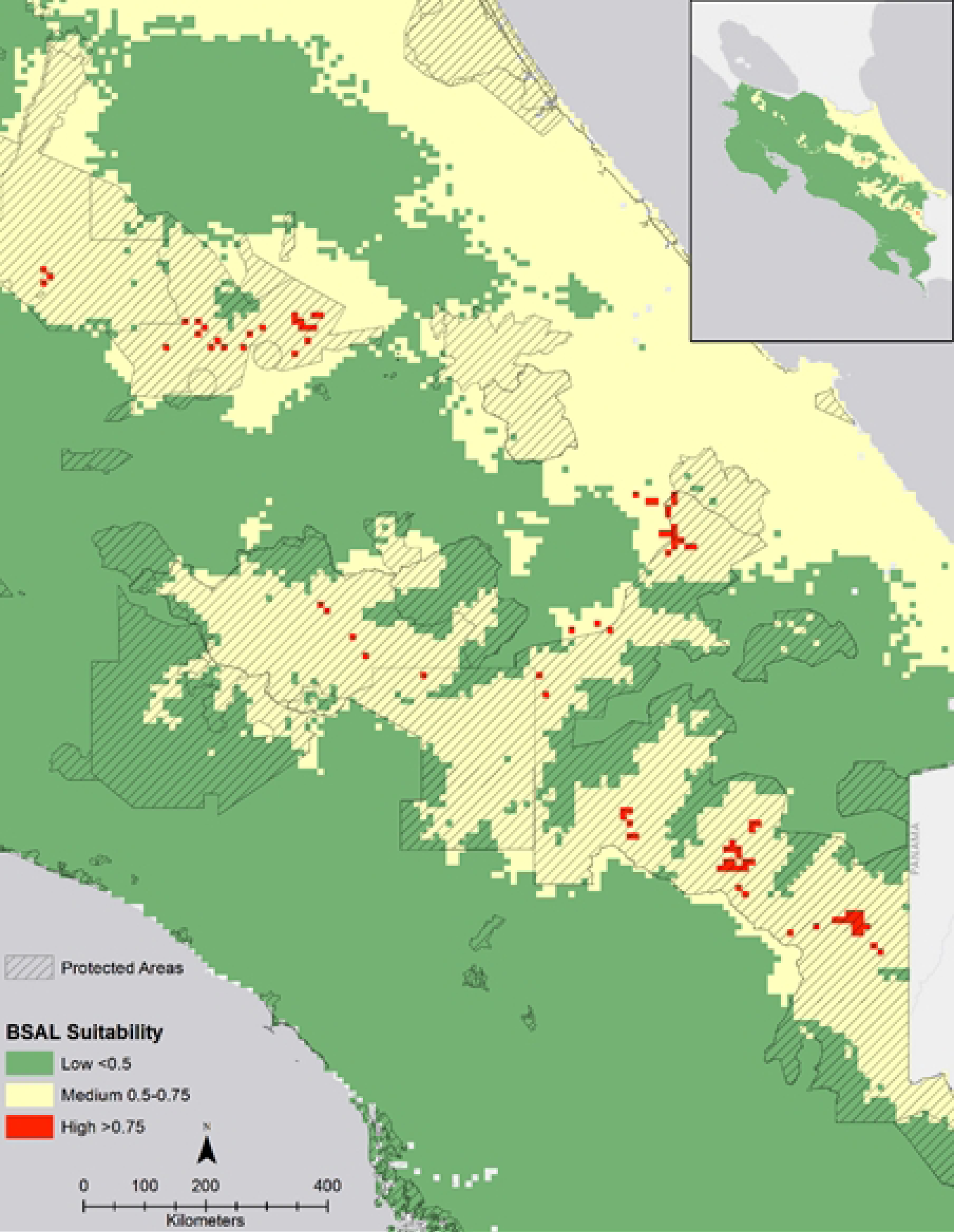
Predicted *Bsal* suitability categorized into high, medium, and low. Protected areas are indicated by black diagonal lines. Insert map shows all of Costa Rica.

### Area highly suitable for Bsal

Eighteen percent of the land modeled to be highly suitable for *Bsal* is inhabited by four or more species of salamanders, and less than 5% of this land contains six or more species. In areas classified as highly suitable, there are a total of 15 species of salamanders, four of which are endangered (Table 1.0). Ninety-two percent of this land falls within protected areas. The protected areas of Parque Internacional La Amistad, Parque Nacional Chirripó, Parque Nacional Tapantí - Macizo Cerro de la Muerte, Parque Nacional Braulio Carrillo, and the protected areas of the Central Volcanic Mountain Range Reserva de la Biosfera Cordillera Volcánica Central all have habitats modeled to be highly suitable (Figure 5). High *Bsal* suitability overlaps with high alpha salamander biodiversity in Reserva de la Biosfera Cordillera Volcánica Central, Parque Nacional Braulio Carrillo, and in Parque Internacional La Amistad along the border of the protected area Río Banano (Figure 6).

**Figure 6.**
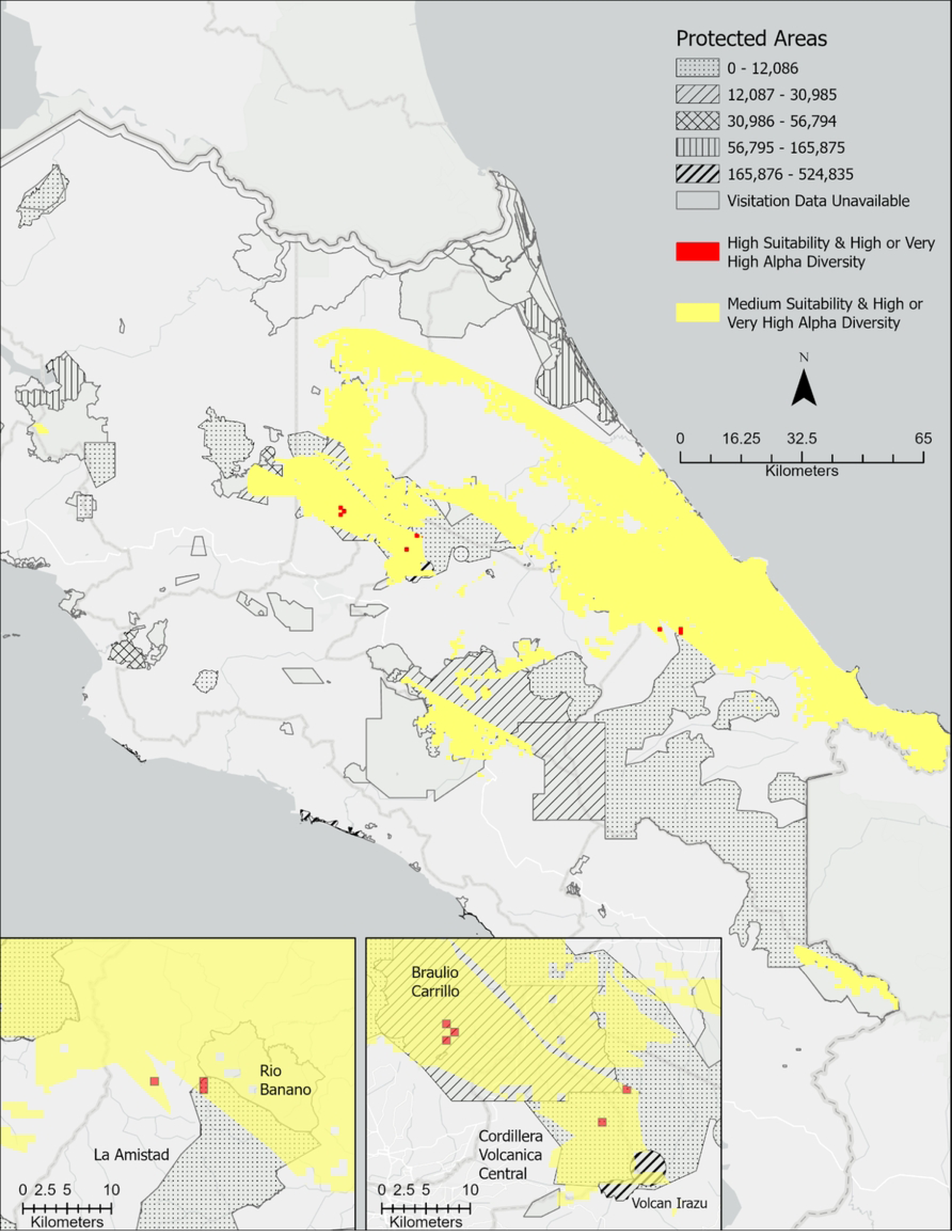
Areas of overlap between predicted moderate and high *Bsal* suitability and high and very high salamander diversity. Areas of both *Bsal* moderate suitability and high/very high salamander diversity are shown in yellow. Areas highly suitable for *Bsal* with high/very high salamander diversity are shown in red. Protected areas are outlined and the number of visitors in 2018 is indicated by the pattern in the legend.

Areas that are highly ecologically suitable for *Bsal* within Parque Nacional Braulio Carrillo, Cordillera Volcánica Central Biological Reserve, and Parque Nacional Tapantí - Macizo Cerro de la Muerte have high RRI values, indicating that the salamander species found within these protected areas are highly endemic to said areas (Figure 7). See Table II Appendix I for these species. In contrast, areas highly suitable for *Bsal* within Parque Nacional Chirripó and Parque Internacional La Amistad overlap with areas that have relatively low RRI levels, indicating the salamander species present in these spaces have a wider distribution across the country.

**Figure 7.**
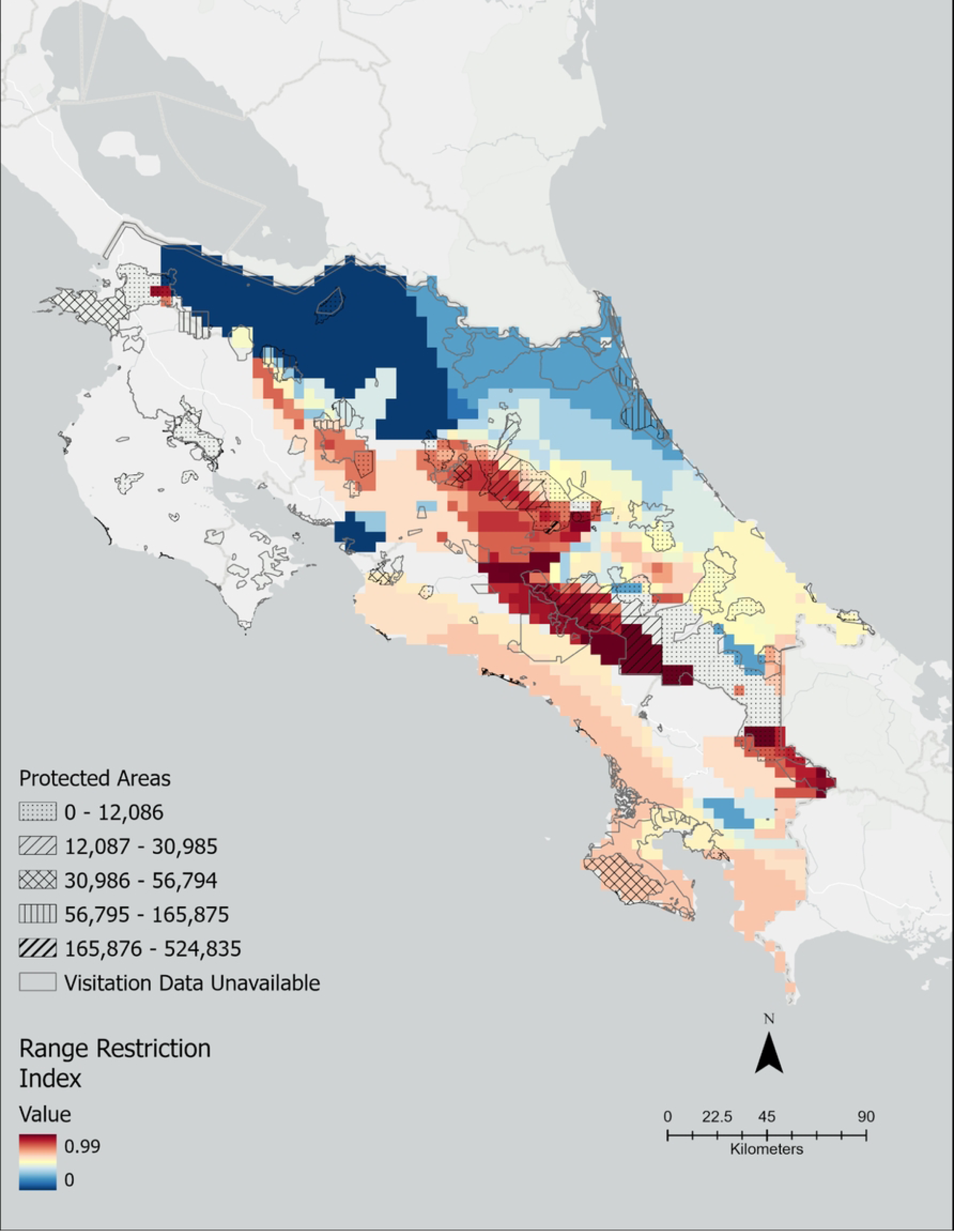
Costa Rican salamander Range Restriction Index results. Range Restriction Index values mapped such that darker red indicates salamander species found within this pixel have a restricted range across Costa Rica. Protected areas are outlined and the number of visitors in 2018 is indicated by pattern.

### Area of moderate suitability

Of the area considered moderately suitable for *Bsal*, 18% contains between six and eight species of salamanders and 43% contain four or five species. In total, 37 of the 44 salamander species included in this study are found in areas of moderate ecological suitability for *Bsal* (Table 1). These 37 species include the critically endangered *Nototriton major* and 13 other salamanders listed as endangered by the IUCN. Just over 56% of land modeled to be moderately suitable for *Bsal* is within protected areas (Table 1.0), including parks with considerable tourist visitation, such as Parque Nacional Volcán Irazú, Parque Nacional Tortuguero, Parque Nacional Cahuita, Parque Nacional Volcán Tenorio and Parque Nacional Volcán Arenal, which all had over 100,000 visitors in 2018 (39).

**Table 1.** Salamander species, total land area, and protected land area in areas of high, medium, and low predicted *Bsal* suitability.

| Suitability | Total land area (km<sup>2</sup>) | Area protected (km<sup>2</sup>) | Area with four or more salamander sp. (km<sup>2</sup>) | Salamander species present<br>(*indicates endangered **indicates critically endangered status in Costa Rica) |
| --- | --- | --- | --- | --- |
| High | 78.62 | 72.70 | 16.90 | *Bolitoglossa alvaradoi* *, *B. colonnea*, *B. diminuta*, *B. epimela*, *B. gracilis*, *B. pesrubra*, *B. robusta*, *B. subpalmata* *, *Oedipina alfaroi*, *O. cyclocauda*, *O. gracilis* *, *O. poelzi* *, *O. uniformis*, *Nototriton* *abscondens*, & *N. richardi* |
| Moderate | 11,585.39 | 6,503.18 | 9,153.83 | *B. alvaradoi* *, *B. bramei* *, *B. cerroensis*, *B. colonnea*, *B. compacta* *, *B. diminuta*, *B. epimela* *, *B. gracilis*, *B. lignicolor*, *B. marmorea* *, *B. minutula* *, *B. nigrescens* *, *B. obscura*, *B. pesrubra*, *B. robusta*, *B. schizodactyla*, *B. sombra*, *B. sooyorum* *, *B. striatula*, *B. subpalmata* *, *B. tica* *, *N. abscondens*, *N. gamezi*, *N. guanacaste*, *N. major* **, *N. picadoi*, *N. richardi*, *O. alfaroi*, *O. carablanca* *, *O. collaris* *, *O. cyclocauda*, *O. gracilis* *, *O. grandis* *, *O. poelzi* *, *O. pseudouniformis* *, & *O. uniformis* |
| Low | 39,440.16 | 6,476.12 | 3,855.80 | *B. alvaradoi* *, *B. bramei*, *B. cerroensis*, *B. colonnea*, *B. compacta* *, *B. epimela*, *B. gomezi*, *B. gracilis*, *B. lignicolor*, *B.* |
|  |  |  |  | *marmorea\**, *B. minutula\**, *B.*<br>*nigrescens\**, *B. obscura*, *B.*<br>*pesrubra*, *B. robusta*, *B.*<br>*schizodactyla*. *B. sombra*, *B.*<br>*sooyorum\**, *B. striatula*, *B.*<br>*subpalmata\**, *B. tica\**, *N.*<br>*abscondens*, *N. gamezi*, *N.*<br>*guanacaste*, *N. major\*\**, *N.*<br>*picadoi*, *N. richardi*, *N. tapanti*, *O.*<br>*alfaroi*, *O. alleni*, *O. altura\**,<br>*O. carablanca\**, *O. collaris*, *O.*<br>*cyclocauda*, *O. gracilis\**, *O.*<br>*grandis\**, *O. pacificensis*, *O.*<br>*paucidentata\**, *O. poelzi\**, *O.*<br>*pseudouniformis\**, *O. savagei*, *O.*<br>*uniformis* |

### Identifying Priority Areas for surveillance and other management actions

To identify parks of priority for *Bsal* surveillance, we first found areas of overlap between high *Bsal* suitability and high/very high salamander diversity (i.e. four or more species). Parque Nacional Braulio Carrillo, Parque Internacional La Amistad, Parque Nacional Tapantí - Macizo Cerro de la Muerte, Parque Nacional Chirripó, Reserva de la Biosfera Cordillera Volcánica Central have habitat that is both predicted to be highly suitable for *Bsal* and contains high/very high salamander diversity. We also considered areas that contain an overlap of high/very high salamander alpha diversity, moderate pathogen habitat suitability, and high human visitation in 2018 (>40,000 visitors). These areas include Parque Nacional Volcán Irazú, one of the most visited parks in 2018 with 400,000 visitors, Parque Nacional Cahuita (∼126,000), as well as Parque Nacional Volcán Poás (∼50,000 visitors) (Figure 8). See Table II Appendix I for more details.

**Figure 8.**
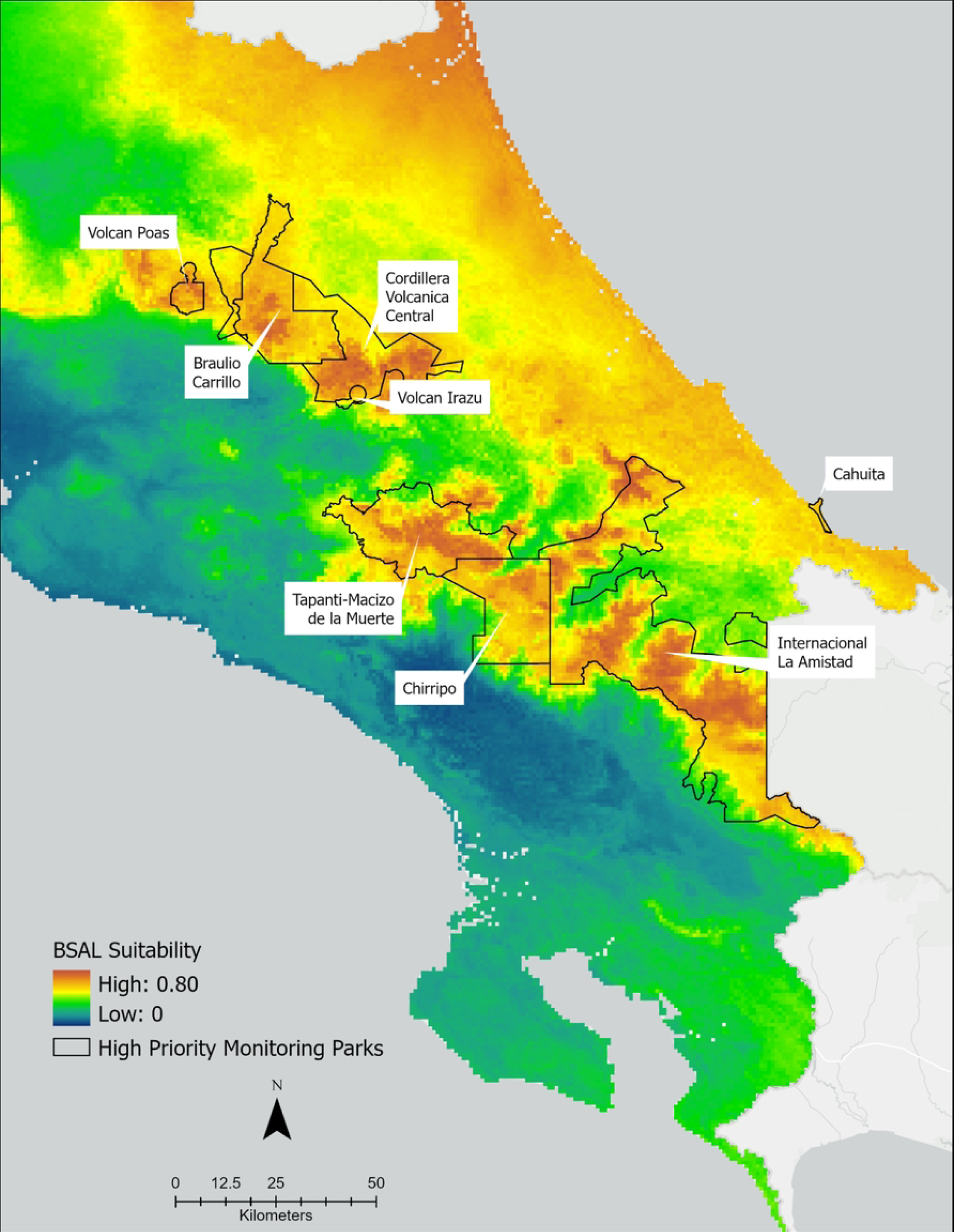
Areas considered high priority for monitoring for *Bsal* introduction. Protected priority areas outlined in black and identified by name overlaid on Ecological Niche Model results for *Bsal* suitability. Areas of warmer colors (reds and oranges) indicate high *Bsal* suitability, whereas cooler colors indicate lower suitability.

## Discussion

*Bsal* is a serious and global threat to urodelans (17–22,59). The pathogen has now been detected in wild amphibians in nine countries and, while it has not yet been detected in the Americas, it is known to infect at least 16 species native to the Americas (60). As our knowledge grows, so does the list of potential host species, with a recent detection in *Rana temporaria* in Germany (61) and a successful experimental inoculation in *Proteus anguinus* (62). We can proactively plan for *Bsal* in part by modeling the potential risk of invasion based on environmental suitability, amphibian biodiversity, and tourist visitation in protected areas. Risk models such as ours provide a rapid and efficient assessment and should be used to provide targeted monitoring and mitigation planning. As *Bsal* has not yet been detected in the Americas, monitoring efforts remain crucially important. Mitigation efforts, such as regulations designed to reduce imports of foreign species, have proven successful in other countries (63).

### Bsal Suitability

Costa Rica’s mountain ranges, specifically the Cordillera Central and Cordillera de Talamanca, are predicted to be the country’s most ecologically suitable areas for *Bsal*. This is in agreement with habitat suitability modeling completed at the scale of the entire American tropics (43). Additionally, suitability is high in the Caribbean tropical forests of northeastern Costa Rica (Figure 4). This was not anticipated because this region’s ambient temperatures frequently range outside of *Bsal’s* ideal thermal niche. Using ecological niche modeling, Basanta et al. (42) found high ecological suitability for *Bsal* in areas of Mexico bioclimatically similar to the northeastern coast of Costa Rica: tropical wet and rainforests with high ambient temperatures and high levels of precipitation. Basanta’s model and our model both found precipitation and temperature to be highly predictive of *Bsal* ecological suitability. It is possible that the high precipitation in the northeast of Costa Rica and the aforementioned regions of Mexico offsets the impact of ambient temperature on *Bsal* suitability in our model. It is also important to consider the role microhabitats play in creating thermal buffers in tropical regions. Tropical microhabitats have been found to be 1-2°C cooler than the macrohabitat’s mean ambient temperature and upwards of 3.5°C cooler than the macrohabitat’s maximum ambient temperature (64,65). Our model relies on remotely-sensed bioclimatic data modeled at 1 km resolution; thus, we are unable to capture any microhabitats important to amphibians. The availability of these microhabitats as potential thermal refuges for *Bsal* as well as its hosts should be considered in the future development of fine scale occupancy models and execution of pathogen surveillance measures.

### Costa Rican Salamander Distribution & Bsal

Over 20% of Costa Rica’s landmass is ecologically suitable for *Bsal*. Though areas identified as highly suitable for *Bsal* are within or adjacent to protected areas and have relatively lower salamander alpha diversity, they could provide important footholds for *Bsal* introduction in Costa Rica. *Bsal* is a density-independent pathogen and thus could spill over into more biodiverse areas even in the absences of highly connected host populations. Additionally, the salamander species in areas highly suitable to *Bsal* have high rates of endemism (Figure 7), presenting not only the risk of severe population decline due to infection and clinical disease, in the event of *Bsal* introduction, but also extirpation.

Our results must be interpreted with some degree of uncertainty, as the susceptibility of specific Costa Rican salamander species to *Bsal* has yet to be determined. Such knowledge would add specificity to targeted surveillance efforts. This highlights the urgent need for susceptibility trials to be conducted with Costa Rican salamanders, especially those belonging to the genus *Bolitoglossa*. With at least 132 species existing only in the American tropics, this is the most diverse *Plethodontidae* genus and accounts for the majority of Costa Rica’s urodelan biodiversity (37,66). *Plethodontidae* species, which includes all Costa Rican urodelans, are known to experience variable susceptibility to the fungus, with some being highly susceptible to infection, clinical disease, and mortality (18,20,67). As such, the large amount of overlap in *Bsal* ecological suitability and vulnerable salamander biodiversity could present a significant risk to the Costa Rican urodelan community. Understanding *Plethodontidae*’s susceptibility to *Bsal* could greatly help predict and formulate management action plans for the fungus not only in Costa Rica but throughout the Americas (43).

Amphibian communities in Costa Rican have already been significantly impacted by *Bd* related epizootics, with some regions experiencing upwards of a 40% decline in species (9). Though *Bd* surveillance in Costa Rican salamanders has been limited, the pathogen is presumed to be endemic within their populations, given the well documented impact of *Bd* on anurans in Costa Rica and *Bolitoglossine* salamanders in Mexico and Guatemala (40,68). Eastern newts co-infected with *Bd* and *Bsal*, showed an increased susceptibility to *Bsal* and a greater downregulated immune response than newts infected with only *Bsal* (69). In the case of *Bsal* introduction to Costa Rica, it is believed that the combined impacts of both chytrid fungi on host populations would be detrimental. Based on the predictions of our ENM and a similar ENM produced by Puschendorf and colleagues, many areas ecologically suitable for *Bsal* in Costa Rica are also ecologically suitable for *Bd* (9). Such overlap exists in regions that have already experienced significant *Bd*-related amphibian population declines, specifically the Monteverde region and the Las Tablas Protected Zone. Monitoring for *Bsal* in these areas, including a handful of our identified areas of priority outlined in Table II, could be crucial to conserving these already degraded amphibian communities.

### Costa Rican Anuran Distribution & Bsal

There is little overlap between land predicted to be highly suitable for *Bsal* and areas of high or very high anuran diversity (Figure I in appendix); most overlap can be found in the protected areas of Parque Internacional La Amistad and Reserva de la Biosfera Cordillera Volcánica Central. Areas predicted to be moderately suitable for *Bsal* that also contain high and very high anuran diversity span the entire eastern coast. Patches of medium suitability and high to very high anuran diversity include Parque Nacional Volcán Arenal, Rincón de la Vieja, and Parque Nacional Volcán Tenorio, all of which had over 50,000 visitors in 2018. Recent research has exemplified the capabilities of anurans to be competent hosts of *Bsal*. The common midwife toad (*Alytes obstetricans*) and *Bombina* spp have been shown to host subclinical *Bsal* infection and can infect urodelan hosts (19). Another anuran of epidemiological significance is the Cuban tree frog (*Osteopilus septentrionalis*). In laboratory infection trials, wild caught Cuban tree frogs collected in Florida, USA, developed clinical chytridiomycosis when exposed to *Bsal* zoospores. Zoospores isolated from diseased Cuban treefrogs were then used to successfully infect healthy eastern newts, exemplifying the frog’s capability to infect naive urodelan hosts (27). This highly adaptive species, which is popular in the international pet trade, has successfully established itself in 14 countries outside its native range (70,71). As such, it could catalyze harmful spillover events and contribute to the global spread of *Bsal*. The Cuban treefrog is considered an introduced species in Costa Rica, however population(s) size and distribution status within Costa Rica remains nebulous and under-examined (70). Close monitoring of Cuban tree frog populations, both wild and captive, and associated surveillance for *Bsal* are highly recommended as this species’ distribution in Costa Rica changes over time. It would also be prudent to investigate the susceptibility of Costa Rican anuran species, targeting those taxonomically related to the Cuban tree frog, such as the veined tree frog (*Trachycephalus typhonius*). This is the only species in the tribe Lophiohylini, (Cuban tree frogs and relatives) that is native to Costa Rica. It is a species of Least Concern according to the IUCN and inhabits much of Costa Rica’s Pacific slope as well as a small portion of the Caribbean slope. Despite its distribution in warmer, lowland habitats, understanding this species’ potential role as a *Bsal* host could help immensely in mitigating future *Bsal* introductions.

### Suggested management actions

*Bsal* has not yet been detected in Costa Rica (37), but regular monitoring is needed to ensure the risk remains as low as possible. Risk models such as ours provide a rapid and efficient assessment and should be regularly employed with the most recent information available in order to target monitoring. Small scale pathogen monitoring has been conducted in Parque Nacional Volcán Poás, Parque Nacional Tapantí - Macizo Cerro de la Muerte, and Parque Internacional La Amistad, three of the priority areas identified in this paper (37). While all animals sampled in this study were negative for *Bsal*, the sample size was small, only encompassing four salamander species, and the continued monitoring of these at-risk populations is essential. Mitigation efforts, such as regulations designed to reduce imports of foreign species, have proven successful in other countries (63). We additionally encourage the development of educational tools designed to inform the public and researchers on amphibian conservation, diseases, and easy actions they can take to limit pathogen transmission. Such actions include responsibly recreating (i.e., limiting recreation to designated trails) and regularly cleaning recreational equipment, especially when visiting protected areas with a high risk of pathogen introduction. Various studies have demonstrated the efficacy of regulatory signage in natural areas in reducing unwanted behaviors, such as off trail hiking and encroaching upon areas of particular ecological value (8,72–74). Similar studies have additionally shown that such signage is particularly impactful when designed with accessibility and approachability in mind: i.e., visually appealing, non verbose, and written with amicable language (72). Other studies have demonstrated the efficacy of household cleaners, such as 4% bleach, in killing both *Bsal* and *Bd* (75), presenting financially accessible measures for the public to control pathogen spread. Such management initiatives could significantly reduce the risk of pathogen introductions going undetected. Ideally, such initiatives would be developed in collaboration with engaged local communities when possible, which would only further conservation management and education efficacy, helping to control spillover events, and aid in global amphibian conservation (76,77).

### Conclusions and limitations

In this study, we used ecological niche modeling, salamander diversity, and tourist visitation data in protected natural areas to identify specific locations, in addition to previously identified geographic regions, worthy of monitoring for the emergence of *Bsal*. Compared to Mexico, a much larger proportion of Costa Rican landmass is predicted to be suitable for *Bsal* (42). We have identified that ∼23% of Costa Rica is moderately to highly suitable for *Bsal* and contains a high amount of salamander diversity. We further pinpointed eight priority regions for surveillance based on the overlapping criteria of *Bsal* ecological suitability, salamander biodiversity, and/or high annual human visitation (Table II, Appendix I). It is our hope that the findings of this study facilitate further pathogen surveillance and prioritized efforts in characterizing the susceptibility of salamander and anuran species within high risk areas in Costa Rica. We additionally encourage the development of educational tools in collaboration with local communities designed to inform the public and researchers on amphibian conservation, diseases, and easy steps they can take to limit pathogen transmission, such as responsible recreation and regularly cleaning of recreational equipment, especially when visiting protected areas with high human visitation.

Remote sensing data and ecological niche models are often crucial tools in biodiversity preservation and conservation work, allowing research at an expansive extent with relatively low financial and personnel cost. Working at such a scale entails generalizations and limitations, particularly when the species of interest is significantly small. Our model does not capture microhabitats, thus future work should be realistic about interpreting this model and cognizantly integrate *in-situ* and local knowledge of the landscape. This risk analysis should supplement existing and future studies and identify novel research questions.

## Acknowledgments

Henry C. Adams was funded by an NSF GRFP #2017239636 and Wildlife Disease Association Challenge Grant. MJG was partially supported by NSF DEB grant 1814520 and USDA NIFA Hatch Project 1012932.

## Supporting Information Captions

**S1 Appendix Table I. Human visitation rate per annum for Costa Rican protected areas in 2018** (1). Note, visitation data was not found for all protected areas. Visitation numbers include both international and local visitors.

**S1 Appendix Table II. Priority regions statistics.** This table lists the eight high priority areas suggested for monitoring. For each priority region, the total area in square kilometers that is considered highly and moderately suitable for *Bsal* is given. The average and range of suitability are listed (one indicates the habitat is highly suitable and zero indicates the habitat is completely unsuitable). Park visitation numbers are from 2018.

**S1 Appendix Figure I. Areas of overlap between predicted moderate and high *Bsal* suitability and high and very high anuran diversity.** Areas of both *Bsal* moderate suitability and high/very high anuran diversity are shown in yellow). Areas highly suitable for *Bsal* with high/very high anuran diversity are shown in red with a close-up shown in the lower, right insert. Protected areas are outlined and the number of visitors in 2018 is indicated by pattern.

**S1 Appendix Figure II. Overlap between Bsal suitability and Veined Tree Frog (Trachycephalus typhonius) range in Costa Rica.** Suitability has been categorized into high (red), medium (yellow), and low (green).

**S1 Appendix Figure III. Salamander gamma diversity in protected and non-protected areas.** Darker shades indicate higher gamma diversity (greater number of endemic species compared to all of Costa Rica). Protected areas are outlined and the number of visitors in 2018 is indicated by pattern.

## Notes

### Competing Interest Statement

The authors have declared no competing interest.

